# DBFE: Distribution-based feature extraction from copy number and structural variants in whole-genome data

**DOI:** 10.1101/2022.02.09.479712

**Authors:** Maciej Piernik, Dariusz Brzezinski, Pawel Sztromwasser, Klaudia Pacewicz, Weronika Majer-Burman, Michal Gniot, Dawid Sielski, Alicja Wozna, Pawel Zawadzki

## Abstract

**Motivation:** Whole-genome sequencing has revolutionized biosciences by providing tools for constructing complete DNA sequences of individuals. With entire genomes at hand, scientists can pinpoint DNA fragments responsible for different cancers and predict patient responses to cancer treatments. However, the sheer volume of whole-genome data makes it difficult to encode the characteristics of genomic variants as features for machine learning algorithms.

**Results:** We present three feature extraction methods that facilitate classifier learning from distributions of genomic variants. The proposed approaches use binning, clustering, and kernel density estimation to produce features that discriminate between two groups of patients. Experiments on genomes of 219 ovarian, 61 lung, and 929 breast cancer patients show that the proposed approaches automatically identify genomic biomarkers associated with cancer subtypes and clinical response to oncological treatment. Finally, we show that the extracted features can be used alongside unsupervised learning methods to analyze genomic samples.

**Availability:** The source code of the presented algorithms and reproducible experimental scripts are available on Github at https://github.com/MNMdiagnostics/dbfe

## 1 Introduction

High-throughput genomic technologies are powerful tools that have dramatically changed the landscape of biological research. In particular, whole-genome sequencing (WGS) is proving to be the tool of the future for pinpointing clinically relevant genetic variation (Dewey *et al*., 2014) and guiding precision medicine (Hou *et al*., 2020). Some of the most interesting genetic traits extracted from whole-genome sequences include copy number variations (CNVs) and structural variants (SVs). These and other genomic aberrations can be used to create predictive models that determine whether a patient will be susceptible to a given treatment (Aung *et al*., 2018; Pilié *et al*., 2019). An example of such a WGS-based predictive algorithm is HRDetect (Davies *et al*., 2017), a classifier designed to predict *BRCA1*/*BRCA2* deficiency and assess the implementation of PARP inhibitor treatment in breast and ovarian cancers (Weil and Chen, 2011).

However, many genomic features are still manually engineered by human experts, hindering the rapid development of WGS-based machine learning models. One reason for this is the sheer volume of variational possibilities in genomic data. For example, it has been shown that CNVs are associated with several rare diseases (Li *et al*., 2020), but such analyses either use simple statistics or require visual inspection of CNV distributions (Requena *et al*., 2021; Collins *et al*., 2020; MacDonald *et al*., 2014). Since there are many types of CNVs and SVs (deletions, inversions, tandem duplications, translocations), manual development of machine learning features requires time-consuming, error-prone, experts’ labor.

Statistical and computer science literature on analyzing distributions includes approaches that verify whether two distributions are statistically significantly different (Massey, 1951; Mann and Whitney, 1947), calculate how much they differ from each other (Vallender, 1974), and visualize those differences (Doksum, 1974; Doksum and Sievers, 1976; Chen and Liu, 2013). Such approaches can be used to determine whether differences between variant distributions of two patient groups exist, but will not automatically transform them into predictive features.

In this paper, we propose three Distribution-Based Feature Extraction (DBFE) approaches for genomic variant lengths. The approaches use quantile binning, clustering, and kernel density estimation, respectively, to transform data easily found in VCF or BEDPE files into interpretable numerical descriptors that can be used to train machine learning models. We discuss the benefits of each approach, analyze their sensitivity to parameter tuning and their interpretability. Experimental results on whole-genome samples derived from patients with ovarian, lung, and breast cancer show that the proposed approaches can automatically create features that differentiate cancer subtypes and treatment response. We also analyze how the generated features can be combined with CNVs of 50 genes based on the PAM50 prognostic test (Parker *et al*., 2009) to achieve better predictive performance. Finally, we assess the explanatory power of the presented approaches by analyzing feature importance and performing an unsupervised learning case study.

## 2 Materials and methods

### 2.1 Data collection and processing

Matched tumor-normal genomes from 219 ovarian, 61 lung, and 929 breast cancer patients, sequenced within three genomic consortia: International Cancer Genome Consortium (https://daco.icgc.org/), The Cancer Genome Atlas (https://portal.gdc.cancer.gov/), Hartwig Medical Foundation (https://www.hartwigmedicalfoundation.nl/data-catalogue/), were downloaded from controlled-access databases after meeting formal criteria (Nik-Zainal et al., 2016; The ICGC/TCGA Pan-Cancer Analysis of Whole Genomes Consortium, 2020). Based on the associated clinical data, ovarian cancer patients were labeled as responders/non-responders to platinum-based or poly (ADP-ribose) polymerase inhibitor (PARPi) therapy, lung cancer patients were labeled as responders/non-responders to immunotherapy (pembrolizumab, nivolumab), and breast cancer samples were labeled according to the cancer’s hormone receptor and HER2 status.

The samples were sequenced using a low PCR amplification or PCR-free library preparation protocols and paired-end 100–150 bp Illumina reads with 350–550 bp insert size. The samples were then analyzed using a stack of open-source software pipelined with Ruffus framework (Goodstadt, 2010). The pipeline started with FASTQ file extraction from the downloaded BAM/CRAM files using SAMtools (Danecek *et al*., 2021) and Picard (Broad Institute, 2019). Next, tumor samples with coverage exceeding 75x were downsampled with Seqtk v1.3-r106 (SeqTk, 2021) to approximately 60x mean coverage. Next, all reads were trimmed using cutadapt v2.10 (Martin, 2011) and mapped to the GRCh37 genome using Sanger’s Cancerit CGPMAP pipeline v3.0.0 (Sanger Institute, 2021). Samples with uniquely mapped read coverage below 20x for either tumor or normal genomes were excluded from the analysis (Cibulskis *et al*., 2013; Alioto *et al*., 2015). After the downsampling stage, the mean tumor samples’ coverage across all datasets, measured using Mosdepth v0.3.2 (https://github.com/brentp/mosdepth), was 48x. Finally, variant calling was performed using Sanger’s Cancerit CGPWGS pipeline v2.0.1 (Sanger Institute, 2021) with somatic CNVs, and SVs identified by ASCAT and BRASS, respectively. Additionally, we used Manta v1.5.0 to detect somatic SVs. As input to the proposed methods, we used CSV files describing the length of individual CNVs/SVs extracted from each sample’s ASCAT/Manta VCF files. However, the proposed approaches and the accompanying software work with any data format convertible into an array of lengths.

### 2.2 Distribution-based feature extraction methods

#### 2.2.1 General problem description

Let *ℒ =* [l_1_, l_2_, …, l_n_] be a vector of multisets of lengths, where each multiset l_i_ = {l_i1,_ l_i2,_ …} represents the lengths of variants in a given sample *i*. Moreover, let *y* = [*y*_1_, *y*_2_, …, *y*_*n*_] be a vector of classes corresponding to each sample. Given ℒ and *y*, the goal of feature extraction is to create a new dataset *X* = *x*_1_, *x*_2_, …, *x*_*n*_, where each example *x*_*i*_ = [*x*_*i*1,_ *x*_*i*2,_ …, *x*_*ip*_] is a representation of sample *i* in which *X*_1,_ *X*_2_, …, *X*_*p*_ correspond to the extracted features (*x*_*i*1_ ∈ *X*_1,_ *x*_*i*2_ ∈ *X*_2,_ …, *x*_*ip*_ ∈ *X*_*p*_).

For example, consider the following list of length sets ℒ = [{1,10,10}, {100,300,200}, {2000,5000}, {7000,8000,10000}] and its corresponding classes *y* = [−, −, +, +]. Based on the length distributions, one could extract features *X*_1_ = ⟦1,300⟧ and *X*_2_ = ⟦2000,10000⟧ that count variants within a range of lengths, and create a dataset as presented in Table 1.

**Table 1.** Sample dataset extracted from sets of lengths ℒ = [{1,10,10}, {100,300,200}, {2000,5000}, {7000,8000,10000}] with classes *y* = [−, −, +, +].

| | $X_1 = [1,300]$ | $X_2 = [2000,10000]$ | $y$ |
| --- | --- | --- | --- |
| $x_1$ | 3 | 0 | – |
| $x_2$ | 3 | 0 | – |
| $x_3$ | 0 | 2 | + |
| $x_4$ | 0 | 3 | + |

Since studies have shown that different lengths of variants accumulate in different types of cancer and that these patterns of accumulation correlate with response to specific treatments (Davies *et al*., 2017), a natural way of achieving the above-described goal is to create features by counting the number of variants in given length ranges. The problem is that these ranges need to be specified separately for each analysis and require expert knowledge, which may be uncertain or unavailable. Therefore, given a desired number of features as a parameter, we aim to provide a tool for automatically identifying such ranges of genomic variant lengths.

The most straightforward method for transforming vectors of variant lengths into a set of features is to divide the range of all possible lengths into a fixed number of bins corresponding to sub-ranges of equal width. After such a transformation, each bin (feature) holds the fraction or the total number of variants of lengths defined by a given sub-range. Although this method is efficient and straightforward to implement, it takes nearly zero information from the dataset into consideration when defining the features. Therefore, such equal-width binning does not reflect the true distribution of lengths in a given dataset. Below we propose three alternative methods to achieve this goal, together with a visualization explaining a given decision.

#### 2.2.2 Genomic variant features based on quantile binning

As a direct alternative to equal-width binning, we propose using quantile binning. In quantile binning, ranges are divided so that each bin contains the same number of variants (Fig. 1, left). In this scenario, the partition is based on the general distribution of all lengths in all samples. As a result, unlike equal-width binning, this approach avoids situations where some bins are empty while others compress all of the information. Nevertheless, this approach still takes the information about the distribution only partially into consideration because it does not actively search for the true underlying distribution of the dataset but only aims at creating equally filled partitions.

**Fig. 1.**
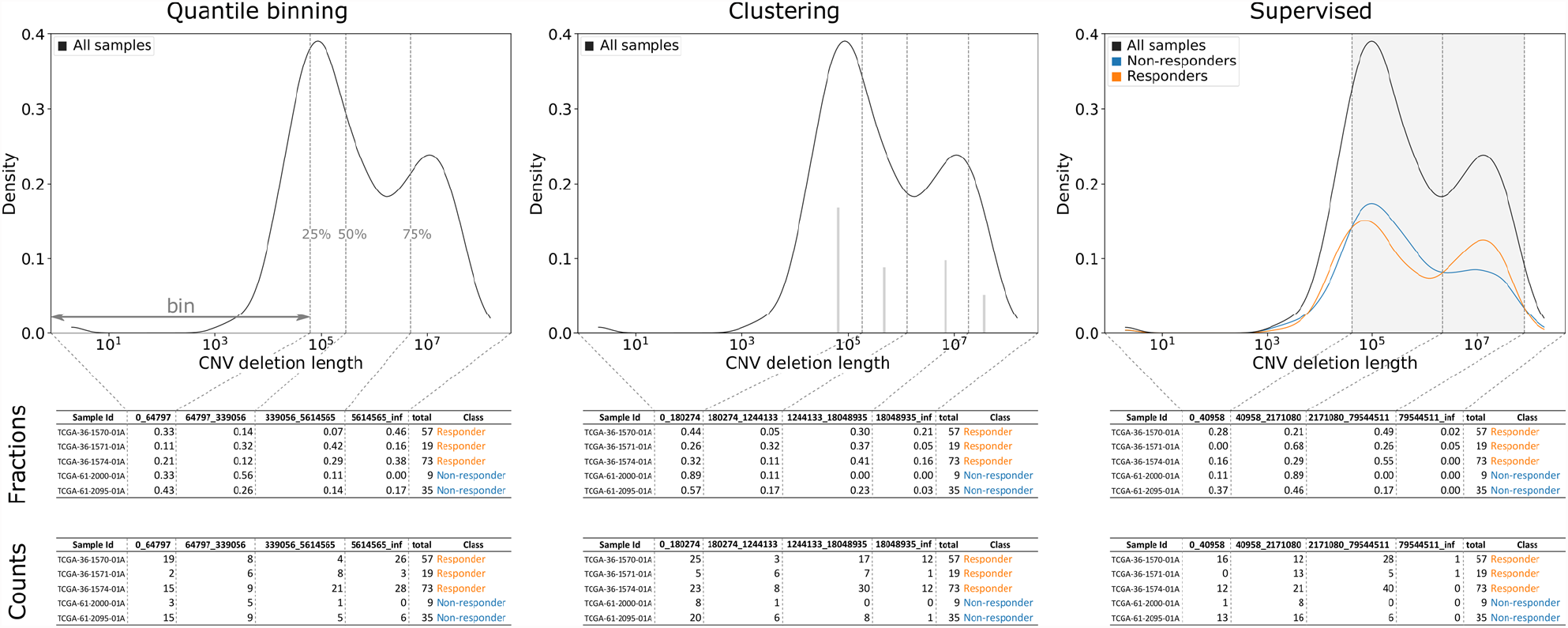
Schematic of quantile binning (left), clustering (middle), and supervised (right) distribution-based feature extraction. Each plot shows the distribution of variant lengths, in this case CNV deletion lengths in ovarian tumors. Note that lengths are presented on a logarithmic scale (x-axis). Vertical dashed lines depict bins, i.e., variant length ranges that define the extracted features. In quantile binning bins are defined by quantiles (dashed gray lines), in clustering by Gaussian cluster centers (solid gray lines), and in the supervised approach by the difference between class-wise distributions, here responders (orange) and non-responders (blue). Based on bin definitions, features can be extracted as fractions of variants within a given range (upper tables) or as counts of variants with the range (bottom tables).

#### 2.2.3 Genomic variant features based on clustering

A more informed approach that tries to detect the distribution underlying the lengths of variants in the dataset is to approximate it with multiple Gaussians. The intuition behind this approach is that if two or more Gaussian distributions can be distinctly detected in the distribution of all lengths in the dataset, they indicate ranges of higher concentrations of variants and therefore are natural candidates for bins. That is why, as a second method for feature extraction, we propose to detect such Gaussians using EM clustering (Moon, 1996), which identifies these regions and automatically assigns each length to an adequate bin based on the given bin’s distribution (Fig. 1, middle). The limitation of this approach is that it requires the number of bins to be defined upfront. Although the same problem concerns quintile binning, here, this decision is more consequential as an incorrectly defined number of bins will force the algorithm to shift from the actual Gaussians, and the benefit of using this method will be diminished.

#### 2.2.4 Genomic variant features based on kernel density estimation

The quantile binning and clustering approaches are applicable to unsupervised scenarios, in which the classes of samples are not used when partitioning lengths. However, when classes are available, one may be interested in defining bins in a way that maximizes the discriminative power of the resulting features. In such cases, we propose an approach based on kernel density estimation (KDE). The method works by:

1. approximating the density of the distribution of lengths separately for samples in each class,
2. identifying the cross points of the two densities,
3. using these points as bin boundaries for features, and
4. optionally filtering out features with low discriminative power.

The right panel of Fig. 1 presents an example of supervised feature extraction, with class-wise length distributions shown with orange and blue lines, filtered regions defined by cross points highlighted by gray areas, and the final bin boundaries depicted by dashed vertical lines.

The intuition behind this approach is that the numbers of variants between two adjacent cross points are, by definition, distinct for different classes. Moreover, the greater the difference between the two densities in a given range, the stronger the discriminative power of the resulting feature. An additional advantage of this method is that it automatically sets an upper bound on the number of generated features, as there is only a finite number of cross points between two distributions. Therefore, the method can automatically detect the maximal number of features to generate without the user specifying it. Moreover, the features can be easily filtered based on the density differences so that only the ones with the highest discriminative power are selected.

The method exposes several parameters which influence its behavior: *bandwidth* – a smoothing parameter of KDE; the higher the value, the smoother the estimation; *log_scale* – whether the estimation should be done using linear or logarithmic scale; *only_peaks* – whether the whole spectrum of lengths should be used when calculating the features, or should only the most discriminative ones be selected; *cv* – whether the density estimation should be done once on all samples or should the process be repeated in cross-validation to account for potential outliers, i.e., singular samples with a significantly higher number of variants; *value_range* – whether the range of lengths should be defined by minimum and maximum value in the dataset or should it be clipped to a specific range. Due to the wide range of lengths of CNVs and SVs, in our experiments, we will work on a log scale and will not limit the range in any way. The impact of the remaining parameters on classifier performance will be discussed in the *Parameter sensitivity* section.

#### 2.2.5 Availability

All of the above-described approaches are available as a python package at https://github.com/MNMdiagnostics/dbfe. The code repository includes installation instructions, short examples, and a Jupyter notebook containing reproducible scripts for the experiments presented in this paper. Since the methods were designed with raw genomic data analyses in mind, the package is equipped with scripts to extract variant lengths from typical VCF files. Moreover, because the method uses the information found in the samples to create feature definitions, it needs to be used only on the training samples to avoid data snooping and potential overfitting. To ensure that such information leakage can be easily avoided, the package is compatible with scikit-learn transformers (Pedregosa *et al*., 2011). Therefore, distribution-based feature extraction can be added as part of a predictive pipeline, together with any scikit-learn estimator or preprocessing method.

### 2.3 Machine learning methods and experimental design

To assess the usefulness of the proposed feature extraction methods, we have tested them against different types of variants in combination with various classifiers on three different classification problems. As input, we tested lengths of six types of variants: three types of CNVs (deletions, diploid copy number, duplications) and three types of SVs (inversions, deletions, tandem duplications). To verify how the extracted features affect different classifiers, we evaluated the performance of four algorithms (Hastie *et al*., 2009; Breiman, 2001; Fix and Hodges, 1989): logistic regression (LR), naïve Bayes (NB), random forest (RF), and *k*-nearest neighbors (kNN). These classifiers were selected because of their complementary nature: logistic regression is a linear classifier, naïve Bayes is a probabilistic classifier, random forest is a tree-based ensemble method, and *k*-nearest neighbors is an instance-based learner. All the classifiers were pipelined with a standard scaler to ensure different feature value ranges did not affect the models.

To evaluate the recognition rate of the trained classification models, the collected datasets were divided into training and holdout testing sets. The holdouts constituted 30% of each dataset and were selected as a random stratified sample before any data preprocessing or analysis. Sensitivity tests and parameter tuning were performed using two repetitions of stratified 5-fold cross-validation (2×5 CV). We evaluated the classifiers using the area under the ROC curve (AUC). Due to the large number of experiments, in the main text we only present the most representative plots and evaluation metrics; the reader is referred to the online supplementary material and code repository for additional results.

## 3 Results

### 3.1 Parameter sensitivity

Using the training portions of the datasets, we verified how the parameters of each DBFE approach affect classifier performance. The first aspect that was checked was whether classifiers perform best using features encoded as length counts, fractions, or both. Stratified 5×2-fold cross-validation results show that using fractions and counts together usually offers better AUC performance than using just one of the representations (Supplementary Fig. S1). The difference is most significant for random forests (RF) and naïve Bayes (NB), which handle larger numbers of features better than *k*-nearest neighbors (kNN) and logistic regression (LR). Therefore, in all the subsequent experiments, we used fractions and counts together.

We also verified how changing the number of features generated by the quantile and clustering approaches affected the classifiers. Here, the best number of features was dataset- and classifier-dependent; LR and kNN worked better with fewer features, whereas RF and NB used more features (Supplementary Fig. S2). In general, when the number of features was too high, the classifiers’ performance, especially LR, degraded. Therefore, when using quantile and clustering DBFE, the number of features needs to be tuned for each dataset.

We also analyzed how parameters *only_peaks, cv*, and *bandwidth* affected classifiers using the supervised approach. Using only features defined by distribution peaks offered slightly better performance for LR but degraded performance for NB, RF, and kNN (Supplementary Fig. S3). Once again, this may be connected to how the different classifiers handle multiple, potentially correlated, features. Using cross-validation (2, 3, 5, and 10-fold) to smooth out distribution peaks did not significantly affect any of the analyzed classifiers (Supplementary Fig. S4). Indeed, we did not find any singular samples with significantly more variants; therefore, such smoothing was unnecessary in this study. Finally, changing the band-width of kernel density estimation (KDE) in the supervised approach did visibly impact classifier performance (Supplementary Fig. S5). Depending on the dataset and variant type, classifier performance either slowly degraded with growing bandwidth or rapidly increased and stabilized. Therefore, the default value of KDE bandwidth was set to 0.5. However, for best performance, KDE bandwidth in the supervised approach should be tuned for each dataset, just like the number of features for the quantile binning and clustering approaches.

### 3.2 Predictive performance of classifiers using DBFE

To evaluate the predictive performance of classifiers using DBFE, we combined features of each variant type within a given approach (quantile, clustering, supervised). The parameters for each approach were chosen based on the results of sensitivity tests. We compared the performance of classifiers using DBFE against those using CNVs of 50 genes based on the PAM50 prognostic test (Parker *et al*., 2009) and those using PAM50 alongside DBFE. Finally, for the ovarian dataset, we contrasted automatically extracted DBFE features with 23 features manually crafted by an expert based on variant length distributions (Supplementary Table S1). The evaluation was performed using the holdout portions of each dataset, which were not considered during training and sensitivity tests.

Looking at the AUC results (Table 2), classifiers using DBFE out-performed those using only PAM50 features on the ovarian, lung, breast ER+/- HER2+ vs. TNBC, and breast ER+ HER2-vs. TNBC datasets. Only on the breast ER+/- HER2+ vs. ER+ HER2-dataset the PAM50 features achieved a consistently better result than DBFE. This is not surprising as the distinction between these subtypes relies on the *ERBB2* gene amplification, which is part of the PAM50 feature set. For all the other datasets, DBFE features either alone or in combination with PAM50 usually offered the best performance. It is also worth noting that the manually engineered features for the ovarian dataset were the best choice for logistic regression, but were worse than DBFE for all the other classifiers.

**Table 2.** The area under the ROC curve (AUC) on holdout data for classifiers using different types of features.

| Dataset | Classifier | AUC |  |  |  |  |  |  |  |
| --- | --- | --- | --- | --- | --- | --- | --- | --- | --- |
|  |  | Human expert | Quantile | Clustering | Supervised | PAM50 | PAM50 + Quantile | PAM50 + Clustering | PAM50 + Supervised |
| ovarian | LR | **0.757** | 0.649 | 0.657 | 0.665 | 0.611 | 0.656 | 0.640 | 0.610 |
| lung |  | - | 0.425 | 0.212 | **0.462** | 0.288 | 0.375 | 0.337 | 0.362 |
| breast<sup>ER+/-HER2+vsER+HER2-</sup> |  | - | 0.920 | 0.948 | 0.953 | 0.929 | 0.977 | **0.980** | 0.979 |
| breast<sup>ER+/-HER2+vsTNBC</sup> |  | - | 0.919 | 0.879 | 0.941 | **0.976** | 0.967 | 0.936 | 0.970 |
| breast<sup>ER+HER2-vsTNBC</sup> |  | - | 0.894 | 0.918 | 0.921 | 0.867 | 0.933 | **0.940** | 0.933 |
| ovarian | NB | 0.595 | 0.644 | 0.659 | **0.743** | 0.536 | 0.630 | 0.681 | 0.699 |
| lung |  | - | 0.450 | 0.562 | 0.425 | 0.525 | 0.562 | **0.587** | 0.450 |
| breast<sup>ER+/-HER2+vsER+HER2-</sup> |  | - | 0.913 | 0.899 | 0.934 | **0.985** | 0.968 | 0.955 | 0.981 |
| breast<sup>ER+/-HER2+vsTNBC</sup> |  | - | 0.866 | 0.889 | 0.928 | 0.964 | 0.888 | 0.944 | **0.972** |
| breast<sup>ER+HER2-vsTNBC</sup> |  | - | 0.592 | 0.740 | 0.857 | 0.793 | 0.639 | 0.750 | **0.858** |
| ovarian | RF | 0.716 | **0.742** | 0.726 | 0.693 | 0.588 | 0.758 | 0.726 | 0.739 |
| lung |  | - | **0.588** | 0.512 | 0.525 | 0.450 | 0.538 | 0.406 | 0.475 |
| breast<sup>ER+/-HER2+vsER+HER2-</sup> |  | - | 0.956 | 0.950 | 0.934 | **1.000** | 0.997 | 0.996 | 0.999 |
| breast<sup>ER+/-HER2+vsTNBC</sup> |  | - | 0.950 | 0.948 | 0.949 | 0.995 | 0.996 | 0.994 | **0.999** |
| breast<sup>ER+HER2-vsTNBC</sup> |  | - | 0.910 | 0.914 | 0.904 | 0.905 | 0.938 | **0.949** | 0.943 |
| ovarian | kNN | 0.641 | 0.706 | **0.750** | 0.725 | 0.532 | 0.718 | **0.750** | 0.679 |
| lung |  | - | 0.550 | **0.650** | 0.400 | 0.575 | 0.425 | 0.569 | 0.600 |
| breast<sup>ER+/-HER2+vsER+HER2-</sup> |  | - | 0.888 | 0.899 | 0.900 | 0.875 | 0.931 | **0.944** | 0.899 |
| breast<sup>ER+/-HER2+vsTNBC</sup> |  | - | 0.915 | 0.916 | 0.935 | 0.865 | 0.965 | 0.936 | **0.968** |
| breast<sup>ER+HER2-vsTNBC</sup> |  | - | 0.812 | 0.847 | 0.877 | 0.815 | 0.851 | **0.903** | 0.859 |

The best values for each dataset-classifier pair are highlighted in bold.

Interestingly, no one classifier consistently worked better than the others on each dataset. For ovarian samples, the best results were obtained by logistic regression (AUC=0.76), for lung cancer by kNN (AUC=0.65), for breast cancer subtype pairs by random forest (AUC=1.00/0.99/0.95). Seeing that different numbers and types of features work better for different classifiers, this shows the high utility of automatically extracted DBFE features compared to manually engineered ones.

Another noteworthy aspect of these experiments is the number of features used by each approach (Supplementary Table S2). The relations between the approaches were consistent between different datasets and classifiers: quantile binning required the most features (median=161), followed by the clustering (median=148) and supervised approach (median=50). The only exception from this rule was the lung dataset, where the supervised approach produced more features than quantile binning and clustering. This is related to the fact that the lung dataset consists of only 61 samples, out of which only 42 were used for training. As a result, kernel density estimation used by the supervised approach was more prone to noise and overfitting. Nevertheless, the supervised approach typically produces significantly fewer features while retaining similar predictive performance as the other DBFE approaches. On the ovarian dataset, the human expert engineered 23 features, which yielded the most straightforward set of features to interpret. However, it is worth noting that some classification problems require more features than others. In such cases, manual engineering might pose a limitation to predictive performance.

### 3.3 Importance of automatically extracted features

To verify how individual features explain the classes in each dataset, we performed feature importance analysis using the ANOVA F-test statistic. The ANOVA F-test offers a classifier-agnostic view of feature importance and was used to see whether quantile, clustering, supervised, or PAM50 features best differentiate between two classes.

The results of importance analysis (Fig. 2) emphasize the descriptive capabilities of DBFE. In particular, the supervised approach produced features that dominated the importance ranking for the ovarian and lung datasets. Moreover, PAM50 features were outside the top 10 ranking in these two datasets. However, some PAM50 genes were significant for breast cancer problems. More precisely, genes *ERBB2* and *GRB7* played a far more important role than the remaining features in differentiating between HER2+ and ER+ HER2-cancer subtypes, which is expected as these genes are colocalized, co-amplified, and directly related to the HER2 receptor (Lamy *et al*., 2011). Interestingly, however, the distinction between ER+/- HER2+ vs TNBC and ER+ HER2-vs TNBC was captured much better by DBFE features. In total, in the top 10 features of all five of the analyzed datasets, only four features (8%) were from the PAM50 list.

**Fig. 2.**
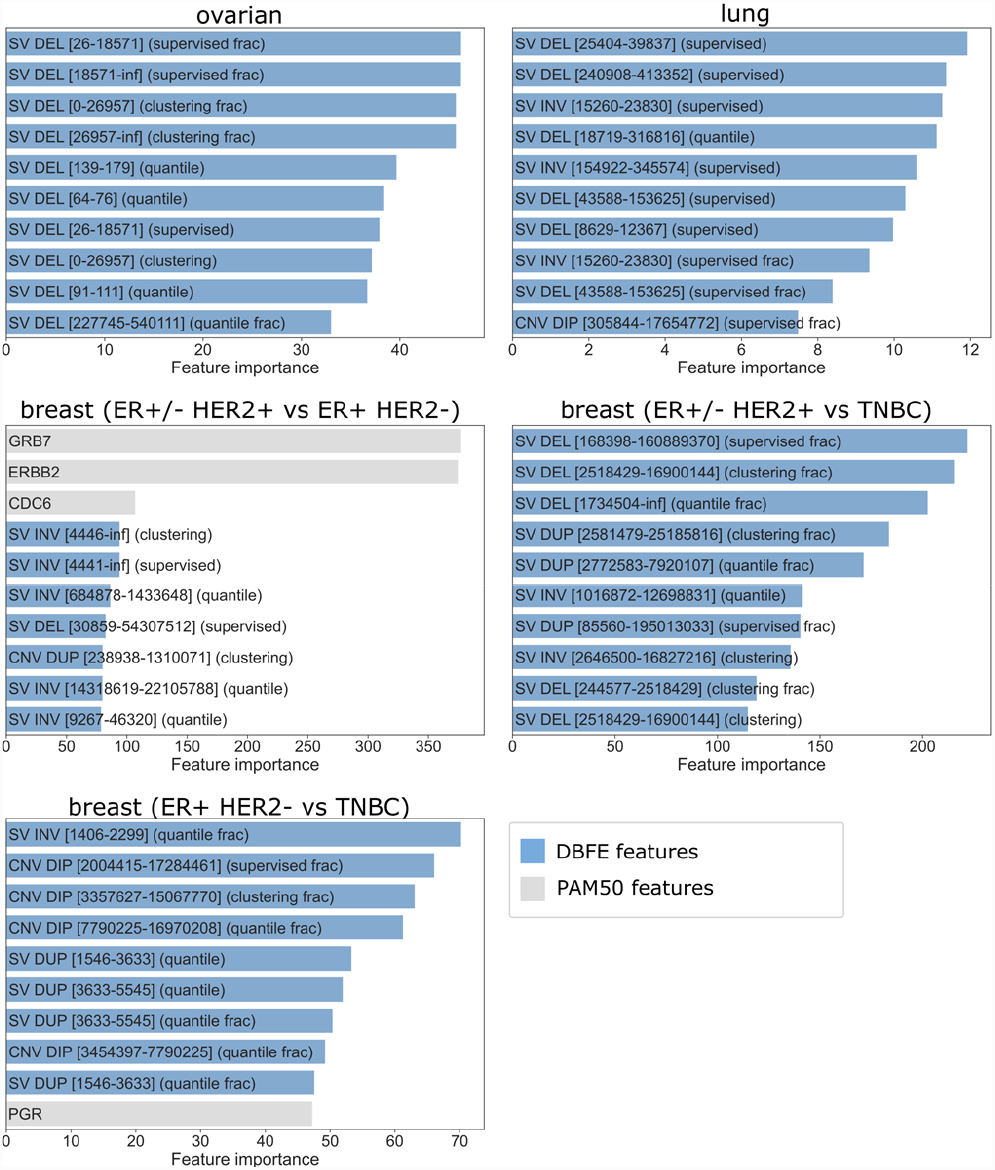
Top 10 most important features for each of the cancer patient datasets. Feature importance calculated using the ANOVA F-test statistic. DBFE features represented by blue bars, PAM50 features represented by gray bars.

### 3.4 Clustering using DBFE

Since the quantile binning and clustering approaches can generate features without class labels, we verified if these approaches can be used to perform interpretable cluster analyses. For this purpose, we used the breast cancer dataset, which consists of 929 samples and would be the most difficult to analyze manually due to its size.

First, we generated four clustering bins for each variant type and produced a total of 54 DBFE features (6 variant types x (4 count features + 4 fraction features + 1 total count)). Then, based on these features, we created a 2D plot using UMAP (McInnes *et al*., 2020) dimensionality reduction (Fig 3A). Interestingly, the grouping of samples in the visualization did not coincide with subtype information available for these samples (Fig 3B). Following a visual analysis of the UMAP plot, we used agglomerative clustering (Hastie *et al*., 2009) to cluster the data into three groups (Fig 3C). Finally, the attribute values of the resulting clusters were visualized by a hierarchical clustering heatmap (Fig. 3D).

**Fig. 3.**
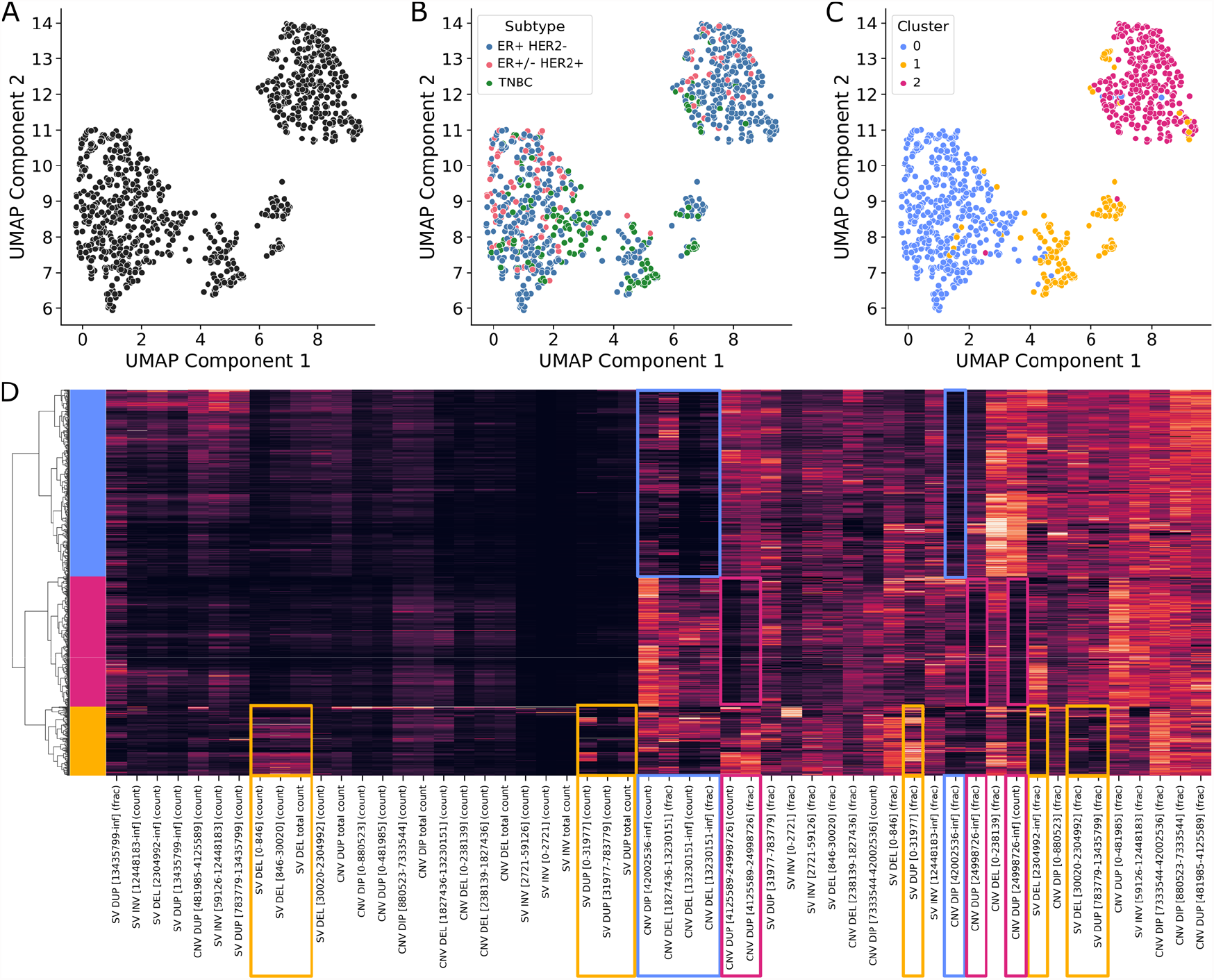
Cluster analysis using features extracted from the breast cancer dataset using the DBFE clustering approach. (**A**) A 2D visualization of the dataset after UMAP dimensionality reduction of the DBFE features. (**B**) The 2D UMAP visualization with points (samples) colored according to the corresponding breast cancer subtype. (**C**) The 2D UMAP visualization with points colored according to clusters assigned using agglomerative clustering with Ward’s linkage method. (**D**) Heatmap presenting the values (tile color) for each DBFE feature (x-axis) in each sample (y-axis). Light tile colors indicate high feature values, whereas dark colors represent low values. The dendrogram on the left presents the clustering of the samples according to the extracted features’ values, with the colors of the leaves representing the same clusters as panel C of this figure.

As already mentioned, the clustering of the samples did not coincide with their assigned subtypes. Almost no HER2-positive samples were classified to Cluster 1, while almost 75% of them fell into Cluster 0. Moreover, the TNBC samples were underrepresented in Cluster 2. The most frequent subtype, ER+ HER2-was evenly distributed between Clusters 0 and 2 but accounts for less than 10% of the samples assigned to Cluster 1. Interestingly, the performed cluster analysis revealed that one of the groups (Cluster 1) consists of samples with a high fraction of short SV deletions and duplications. Further investigation of these samples revealed that over 91% of them are homologous recombination deficient (detected by HRDetect (Davies *et al*., 2017)). Moreover, considering the molecular profile of tumors represented by Cluster 1, almost 55% of the samples harbor *BRCA1* or *BRCA2* loss-of-function mutation. In turn, the second large group of HRD breast cancer tumors (∼19%) has been included in Cluster 0. However, only in 8% of them pathogenic variants within *BRCA1* or *BRCA2* genes have been observed, which may indicate a different HRD origin. Therefore, the clustering helped reveal an alternative, in this case HRD-oriented, way of looking at the samples. Similar results were obtained using features extracted using the DBFE quantile approach (Supplementary Fig. S6).

## 4 Discussion

Distribution-Based Feature Extraction (DBFE) is a set of approaches that automatically generate machine learning features from sets of lengths. DBFE has been tailored particularly for genomic variants (CNVs and SVs) from whole-genome sequencing. Our study shows that variant length features automatically extracted using DBFE can outperform, in terms of predictive performance, PAM50 gene CNVs and features engineered manually by human experts. We have also seen that it is often beneficial to combine gene copy numbers and DBFE features, as they provide complementary information about cancer genomes. A case study on using DBFE to cluster genomic samples (Fig. 3) has also shown the method’s utility in exploratory and unsupervised machine learning settings.

From a biological perspective, experiments with DBFE have confirmed known findings as well as shed light on potentially new avenues for research. For example, structural variants are well-known biomarkers of tumor HRD; therefore, DBFE features were able to help predict patient response to PARPi or platinum-based chemotherapy. On the other hand, CNVs and SVs are not identified as prognostic or predictive features in subtyping, although breast cancer genomes are known to have more structural variants than other tumors. Nevertheless, our study has shown that classifiers using structural features automatically extracted using DBFE were capable of outperforming well-known genes in the PAM50 set. Future studies integrating genomic variability across breast cancer subtypes may help improve the selection of treatments for these patients and help understand the tremendous heterogeneity in breast cancer.

The performance of machine learning approaches like DBFE depends on the number of available training examples. Indeed, the lung cancer dataset results have shown that the supervised DBFE approach, which relies on kernel density estimation, may be susceptible to outliers and noise if the training dataset is too small. The quantile binning and clustering approaches are a better choice in such cases.

One of the benefits of using DBFE is the visual nature of the method. The accompanying library comes with a graphing method that produces density plots similar to those presented in Fig. 1. Such plots facilitate understanding the underlying biological phenomena, which differentiate, e.g., between cancer patients that are drug-sensitive or resistant. Such insights are often as valuable as achieving high predictive performance.

It is also worth contrasting DBFE with deep learning approaches since the recent advancements in this field have made it very popular in genomics (Koumakis, 2020). Deep learning approaches require large amounts of training samples, which is why they are most commonly applied only on readily available types of data, such as whole-exome sequencing, NGS panels, RNA-seq, or miRNA-seq. Such approaches are much rarer in whole-genome sequencing data analyses, where they are mainly used for population-wide studies (Liu *et al*., 2022). When clinical information is required, e.g., in drug response prediction, deep learning approaches are likely to be inadequate because such information is scarce. On the other hand, DBFE works well even with small amounts of data due to the use of global information (length distribution aggregated across all samples throughout the whole genome). Moreover, thanks to its straight-forward feature interpretability enhanced with the provided visualizations, DBFE facilitates the explainability of the resulting models. Such interpretability is paramount in clinical applications yet difficult to achieve with current deep learning models.

The method presented here can also be applied to other variant types, such as small insertions and deletions (indels). In such a case, one would possibly need to limit the range of lengths taken into account and analyze distributions on a linear rather than a logarithmic scale. In addition, the findings from this study could be further advanced by considering the possibility of using the discovered distributions to generate synthetic data. With generative models becoming tools for data imputation, data augmentation, and self-learning (Yelmen *et al*., 2021), automatic extraction of variant distributions could serve as additional genome-wide information for such methods. Hence, the principles of DBFE could serve both as a method for feature extraction and smart data generation.

## Acknowledgments

We thank the international genomic consortia, their researchers, and public funding institutions, who have generously donated the genomic data and clinical metadata for this project. This publication and the underlying study have been made possible by the Hartwig Medical Foundation and the Center of Personalized Cancer Treatment (CPCT) data. This work was also supported by data obtained from ICGC Breast Cancer Working Group and The Cancer Genome Atlas.

## Funding

MP and DB acknowledge the support of PUT’s Institute of Computing Science Statutory Funds.

### Conflict of Interest

none declared.

## Notes

https://github.com/MNMdiagnostics/dbfe

### Competing Interest Statement

The authors have declared no competing interest.

