## Supplement for "DBFE: Distribution-based feature extraction from copy number and structural variants in whole-genome data"

*Genome analysis*

[1]Institute of Computing Science, Faculty of Computing and Telecommunications, Poznan University of Technology, Piotrowo 2, 60-965 Poznan, Poland, [2]MNM Bioscience Inc.,1 Broadway, Cambridge, MA 02142, [3]Institute of Bioorganic Chemistry of the Polish Academy of Sciences, Zygmunta Noskowskiego 12/14, 61-704 Poznan, Poland, [4]Department of Hematology and Bone Marrow Transplantation, Poznan University of Medical Sciences, Poznan, Poland, [5]Faculty of Physics, Adam Mickiewicz University, Uniwersytetu Poznanskiego 2, 61-614 Poznan, Poland

*To whom correspondence should be addressed.

### List of supplementary tables:

- **Table S1**. Features engineered by human expert for the ovarian cancer dataset.
- **Table S2**. Sensitivity analysis (quantile, clustering, supervised): variant fractions and counts.

### List of supplementary figures:

- **Fig S1**. Sensitivity analysis (quantile, clustering, supervised): variant fractions and counts.
- **Fig S2**. Sensitivity analysis (quantile, clustering): number of features.
- **Fig S3**. Sensitivity analysis (supervised): density peaks.
- **Fig S4**. Sensitivity analysis (supervised): cross-validation smoothing.
- **Fig S5**. Sensitivity analysis (supervised): KDE bandwidth.
- **Fig S6**. Cluster analysis using DBFE quantile features.

**Table S1.** Features engineered by human expert for the ovarian cancer dataset.

| Dataset | Variant group | Variant type | Bin ranges | | | | | |
|---|---|---|---|---|---|---|---|---|
| | CNV | DEL | 0-30 | 30-3,500 | 3,500-100,000 | 100,000-Inf | | |
| | | DIP | 0-30 | 30-500 | 500-100 000 | 100,000-Inf | | |
| ovarian | | DUP | 0-30 | 30-100 | 100-350 | 350-3,000 | 3,000-25,000 | 25,000-Inf |
| | SV | DEL | 0-30 | 30-3,500 | 3,500-100,000 | 100,000-Inf | | |
| | | DUP | 0-30 | 30-100 | 100-1,000 | 1,000-30,000 | 30,000-Inf | |

CNV: copy number variation; SV: structural variant; DEL: deletion; DIP: diploid copy number; DUP: duplication.

**Table S2.** Number of features used by each classification approach on each dataset.

| Dataset | Classifier | Number of features | | | | | | | |
|---|---|---|---|---|---|---|---|---|---|
| | | Human expert | Quantile | Clustering | Supervised | PAM50 | PAM50 + Quantile | PAM50 + Clustering | PAM50 + Supervised |
| ovarian | LR | 23 | 136 | 138 | 58 | 50 | 186 | 188 | 108 |
| lung | | - | 76 | 152 | 90 | 50 | 126 | 202 | 140 |
| breast$^{HER2+vsER+HER2-}$ | | - | 174 | 110 | 44 | 50 | 224 | 160 | 94 |
| breast$^{HER2+vsTNBC}$ | | - | 144 | 148 | 50 | 50 | 194 | 198 | 100 |
| breast$^{ER+HER2-vsTNBC}$ | | - | 272 | 184 | 78 | 50 | 322 | 234 | 128 |
| ovarian | NB | 23 | 100 | 84 | 50 | 50 | 150 | 134 | 100 |
| lung | | - | 54 | 86 | 96 | 50 | 104 | 136 | 146 |
| breast$^{HER2+vsER+HER2-}$ | | - | 154 | 170 | 46 | 50 | 204 | 220 | 96 |
| breast$^{HER2+vsTNBC}$ | | - | 168 | 148 | 46 | 50 | 218 | 198 | 96 |
| breast$^{ER+HER2-vsTNBC}$ | | - | 322 | 208 | 48 | 50 | 372 | 258 | 98 |
| ovarian | RF | 23 | 282 | 136 | 64 | 50 | 332 | 186 | 114 |
| lung | | - | 36 | 24 | 140 | 50 | 86 | 74 | 190 |
| breast$^{HER2+vsER+HER2-}$ | | - | 252 | 228 | 78 | 50 | 302 | 278 | 128 |
| breast$^{HER2+vsTNBC}$ | | - | 168 | 218 | 48 | 50 | 218 | 268 | 98 |
| breast$^{ER+HER2-vsTNBC}$ | | - | 272 | 178 | 50 | 50 | 322 | 228 | 100 |
| ovarian | kNN | 23 | 146 | 176 | 44 | 50 | 196 | 226 | 94 |
| lung | | - | 40 | 118 | 70 | 50 | 90 | 168 | 120 |
| breast$^{HER2+vsER+HER2-}$ | | - | 88 | 66 | 42 | 50 | 138 | 116 | 92 |
| breast$^{HER2+vsTNBC}$ | | - | 258 | 124 | 46 | 50 | 308 | 174 | 96 |
| breast$^{ER+HER2-vsTNBC}$ | | - | 244 | 274 | 54 | 50 | 294 | 324 | 104 |
| **Median number of features** | | **23** | **161** | **148** | **50** | **50** | **211** | **198** | **100** |

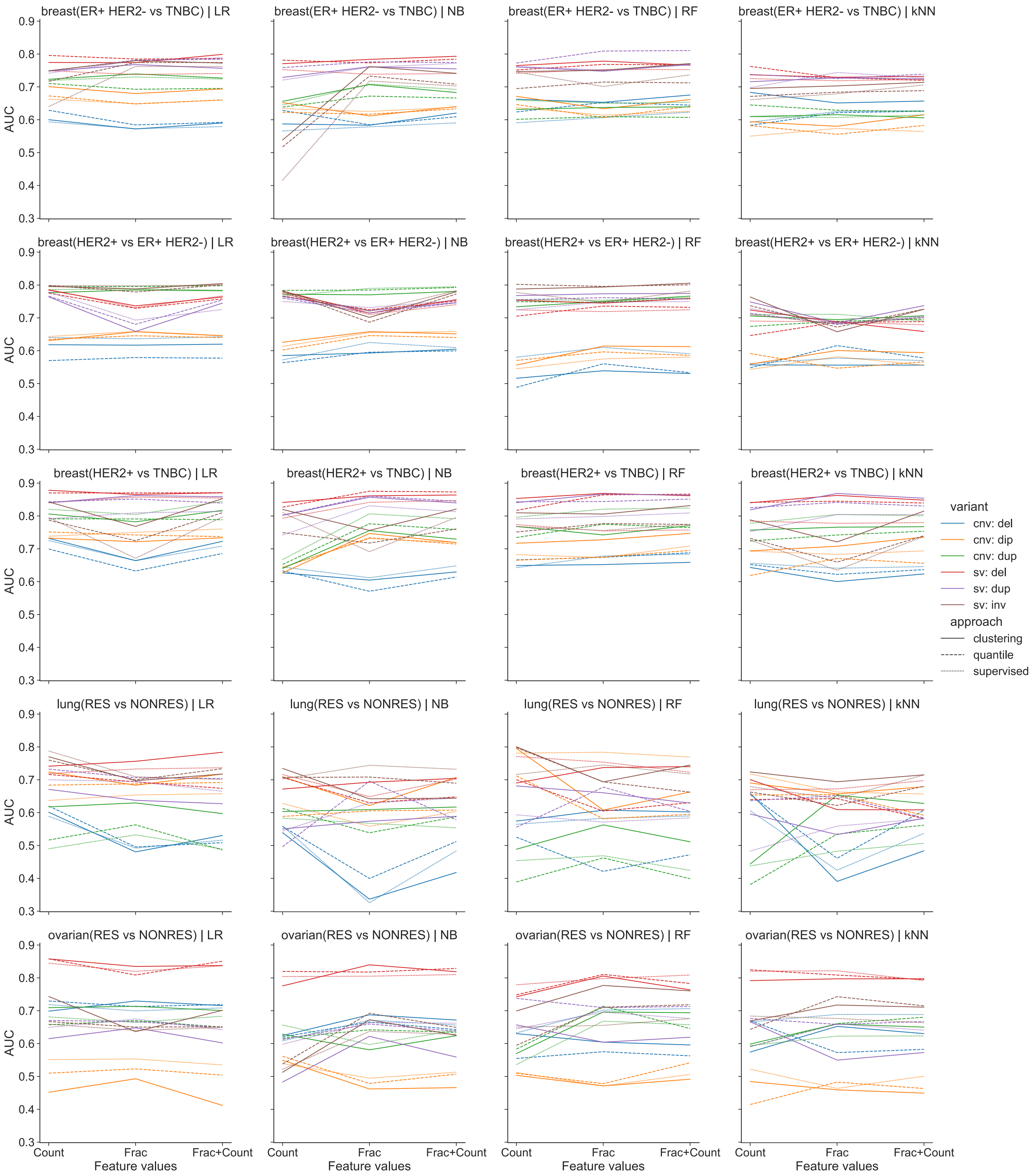


**Fig. S1.** Predictive AUC performance (y-axis) of classifiers depending on whether they used features encoded as length counts, fractions, or both side-by-side (x-axis). Experiments for four classifiers (columns) using three DBFE approaches (line type) to extract features from different variant types (line color) on different datasets (rows). LR: Logistic Regression; NB: Naïve Bayes; RF: Random Forest; kNN: k-Nearest Neighbors.

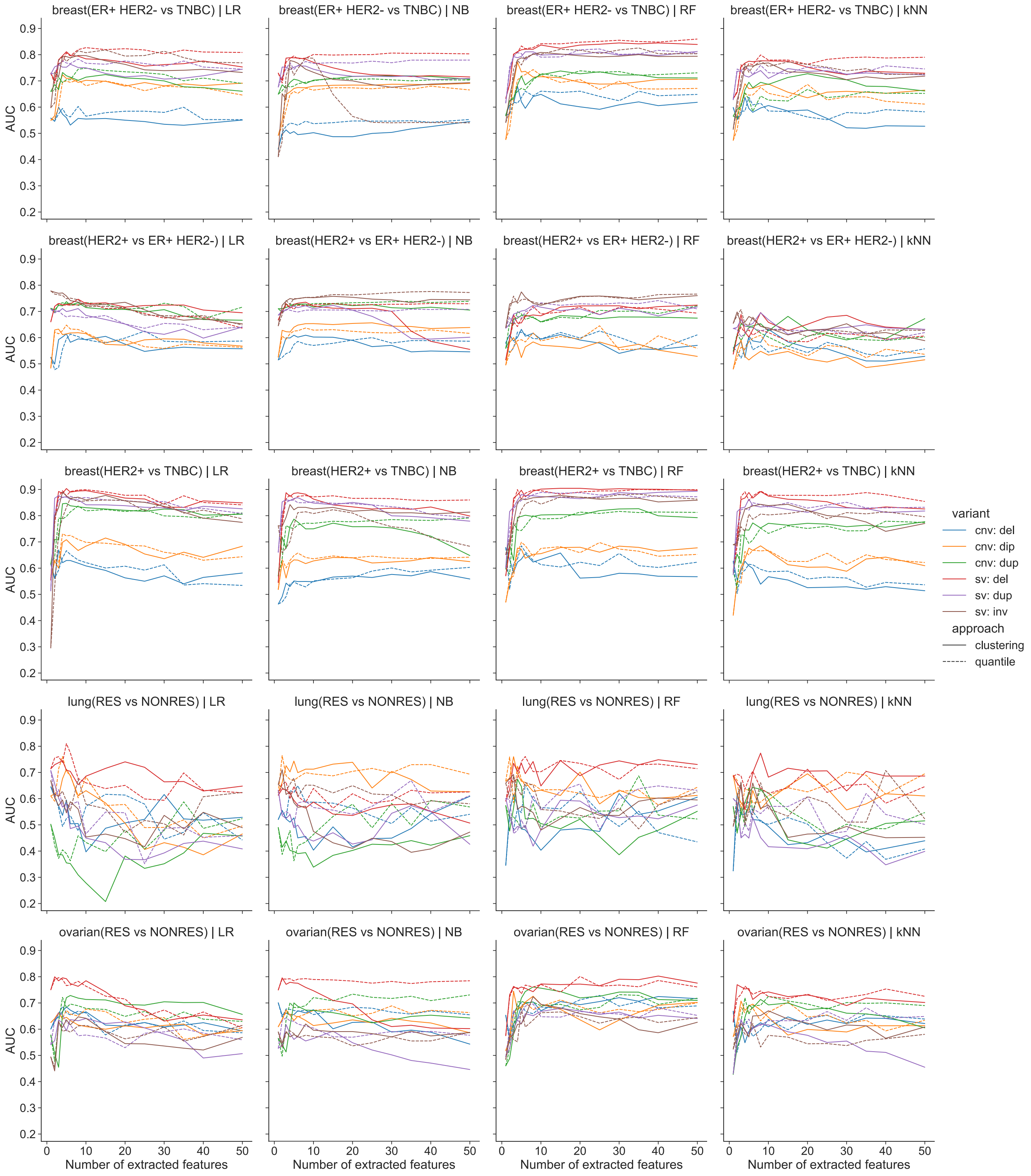


**Fig. S2.** Predictive performance (y-axis) of classifiers depending on the number of extracted features (x-axis). Experiments for four classifiers (columns) using the quantile and clustering DBFE approaches (line type) to extract features from different variant types (line color) on different datasets (rows). LR: Logistic Regression; NB: Naïve Bayes; RF: Random Forest; kNN: k-Nearest Neighbors.

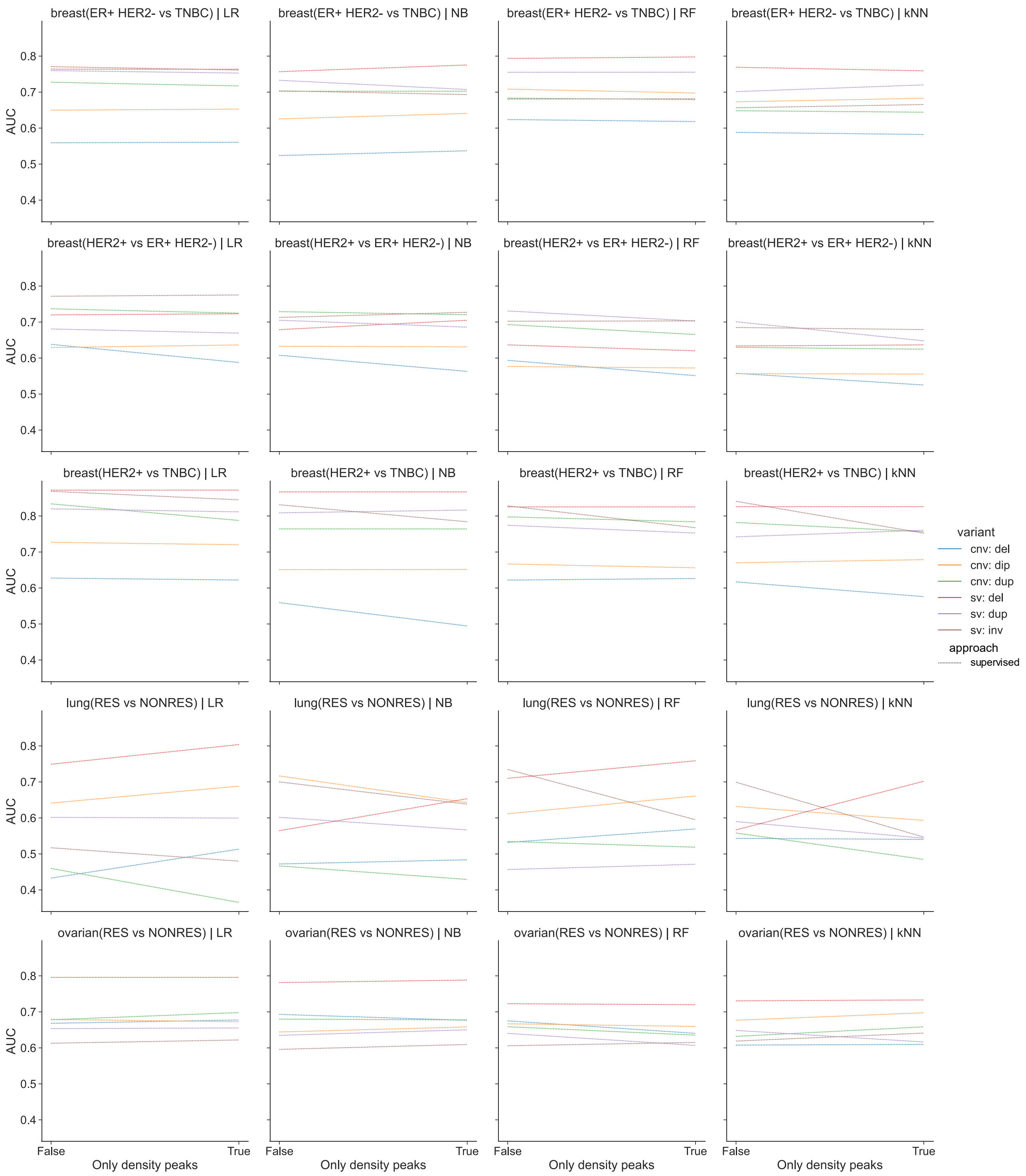


**Fig. S3.** Predictive performance (y-axis) of classifiers depending on whether only density peak regions or entire ranges of lengths were used (x-axis). Experiments for four classifiers (columns) using the supervised DBFE approach to extract features from different variant types (line color) on different datasets (rows). LR: Logistic Regression; NB: Naïve Bayes; RF: Random Forest; kNN: k-Nearest Neighbors.

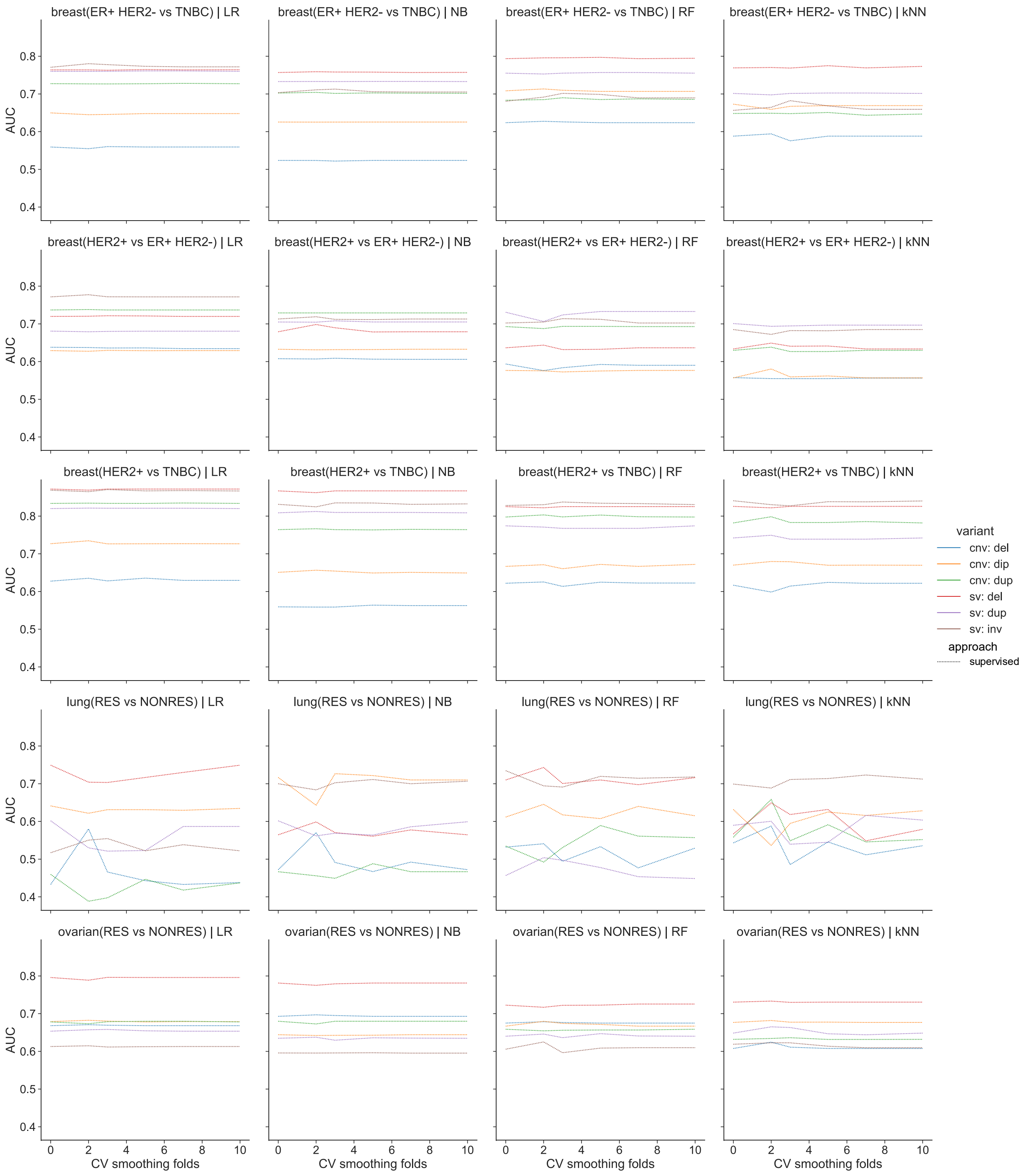


**Fig. S4.** Predictive performance (y-axis) of classifiers depending on the number cross-validation folds used for smoothing (x-axis). Experiments for four classifiers (columns) using the supervised DBFE approach to extract features from different variant types (line color) on different datasets (rows). LR: Logistic Regression; NB: Naïve Bayes; RF: Random Forest; kNN: k-Nearest Neighbors.

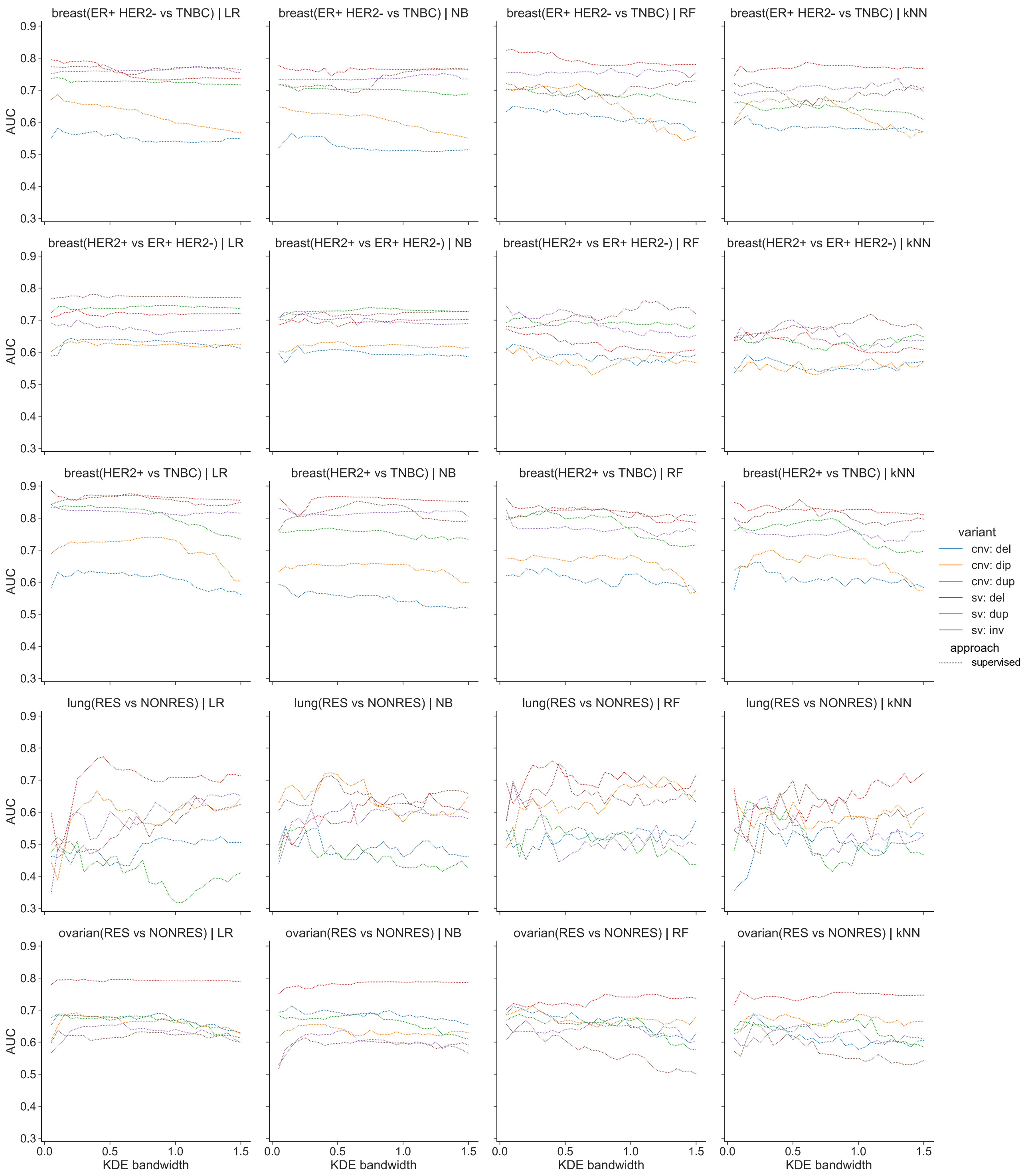


**Fig. S5.** Predictive performance (y-axis) of classifiers depending on the kernel density estimation bandwidth (x-axis). Experiments for four classifiers (columns) using the supervised DBFE approach to extract features from different variant types (line color) on different datasets (rows). LR: Logistic Regression; NB: Naïve Bayes; RF: Random Forest; kNN: k-Nearest Neighbors.

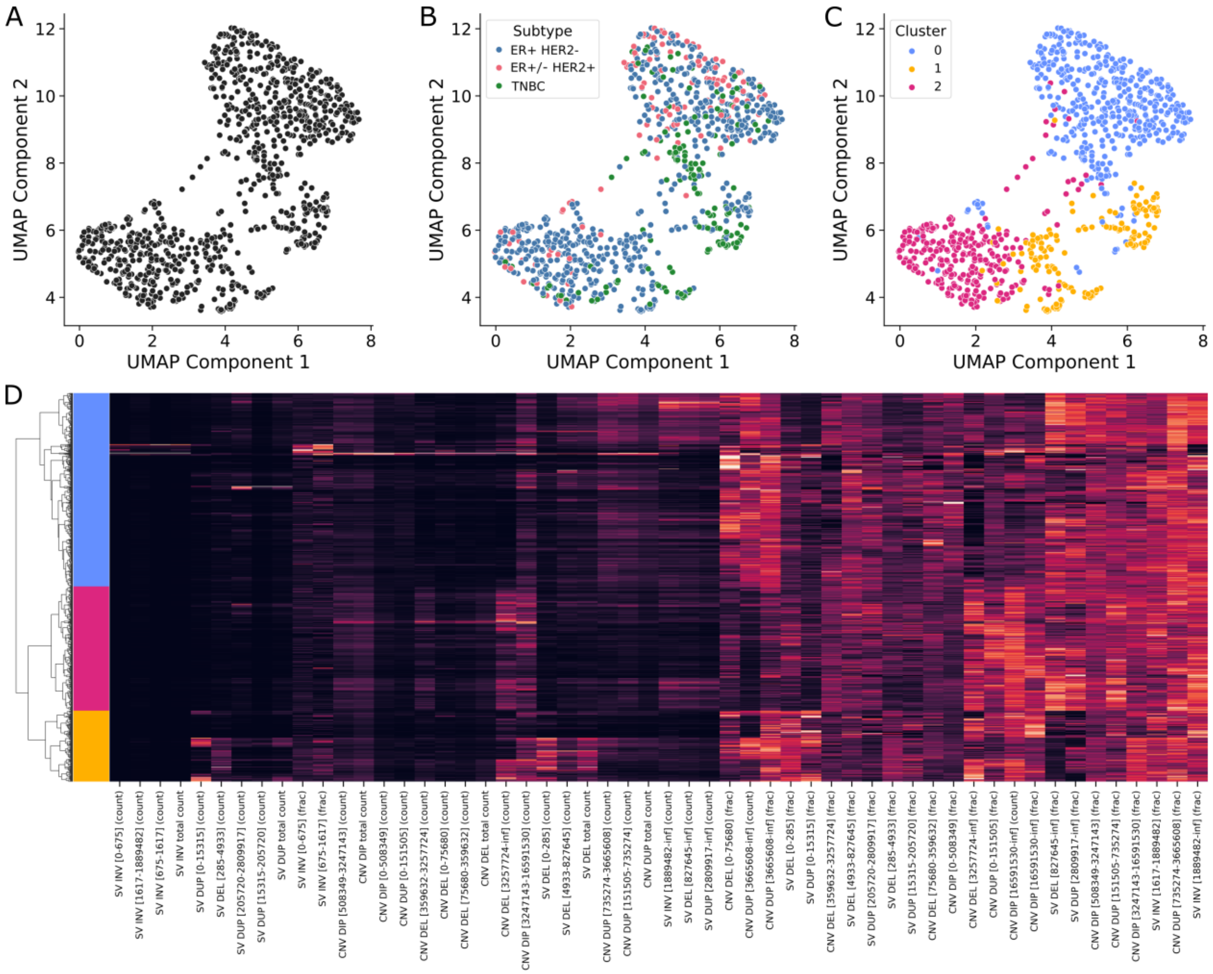


**Fig. S6.** Cluster analysis using features extracted from the breast cancer dataset using the DBFE quantile approach. (**A**) A 2D visualization of the dataset after UMAP dimensionality reduction of the DBFE features. (**B**) The 2D UMAP visualization with points (samples) colored according to the corresponding breast cancer subtype. (**C**) The 2D UMAP visualization with points colored according to clusters assigned using agglomerative clustering with Ward's linkage method. (**D**) Heatmap presenting the values (tile color) for each DBFE feature (x-axis) in each sample (y-axis). Light tile colors indicate high feature values, whereas dark colors represent low values. The dendrogram on the left presents the clustering of the samples according to the extracted features' values, with the colors of the leaves representing the same clusters as panel C of this figure.
